# Haplotype-resolved powdery mildew resistance loci reveal the impact of heterozygous structural variation on NLR genes in *Muscadinia rotundifolia*

**DOI:** 10.1101/2021.11.23.469029

**Authors:** Mélanie Massonnet, Amanda M. Vondras, Noé Cochetel, Summaira Riaz, Dániel Pap, Andrea Minio, Rosa Figueroa-Balderas, M. Andrew Walker, Dario Cantu

## Abstract

*Muscadinia rotundifolia* cv. Trayshed is a valuable source of resistance to grape powdery mildew. It carries two powdery mildew resistance-associated genetic loci, *Run1*.*2* on chromosome 12 and *Run2*.*2* on chromosome 18. In this study, we identified, phased, and reconstructed the two haplotypes of each resistance-associated locus. Haplotype phasing allowed the identification of several structural variation events between haplotypes of both loci. Combined with manual refinement of the gene models, we found that the heterozygous structural variants affected the gene content, with some resulting in duplicated or hemizygous nucleotide-binding leucine-rich repeat (NLR) genes. Structural variations also impacted the domain composition of some NLRs. These findings emphasize the need of generating haplotype-resolved sequences instead of using consensus sequences for identifying haplotype-specific candidate genes. Comparison of the NLRs in the *Run1*.*2* and *Run2*.*2* loci indicated that the two loci are composed of a different number and classes of NLR genes. We provide a list of candidate NLR genes from the *Run1*.*2b* and *Run2*.*2* loci, whose expression suggests a role in powdery mildew resistance in Trayshed. These first complete and haplotype-resolved resistance-associated loci, and their candidate NLR genes, represent new resources to develop powdery mildew-resistant grape cultivars.

## Introduction

Grapevine powdery mildew (PM) is a devastating fungal disease caused by *Erysiphe necator* Schwein. (syn. *Uncinula necator*), an obligate biotrophic ascomycete that can infect all green organs of a grapevine (Gadoury *et al*., 2012). Cultivated grapevines that belong to *Vitis vinifera* (ssp. *vinifera*) are highly susceptible to PM. Fungicide sprays are applied prophylactically to control the disease but are costly (Sambucci *et al*., 2019). Natural resistance to PM exists in several wild grapes. Thirteen PM resistance-associated loci were identified in the last two decades (Dry *et al*., 2019; Karn *et al*., 2021). *Vitis* includes several PM-resistant species, including *V. romanetii* (Ramming *et al*., 2010; Riaz *et al*., 2011) and *V. piasezkii* (Pap *et al*., 2016), which are native to China, *V. vinifera* ssp. *sylvestris* from Central Asia (Riaz *et al*., 2020), the North American *V. cinerea* (Dalbó *et al*., 2001), and the muscadine grape, *Muscadinia rotundifolia* (Pauquet *et al*., 2001; Feechan *et al*., 2013; Riaz *et al*., 2011).

*Muscadinia rotundifolia* is closely related to *Vitis* (Small, 1913) and is resistant to several diseases in addition to PM (Olmo, 1971; Olmo, 1986), including downy mildew, Pierce’s disease, and phylloxera. Two major loci associated with PM resistance were found in *M. rotundifolia. Resistance to Uncinula necator 1* (*Run1*), located on chromosome 12, and its alternative form, *Run1*.*2*, were identified in *M. rotundifolia* G52 and Trayshed, respectively (Pauquet *et al*. 2001; Riaz *et al*., 2011). *Run2*.*1* and *Run2*.*2* were identified on chromosome 18 of *M. rotundifolia* Magnolia and Trayshed, respectively (Riaz *et al*., 2011). Both haplotypes of Trayshed’s *Run1*.*2* were associated with PM resistance and designated *Run1*.*2a* and *Run1*.*2b* (Feechan *et al*., 2015).

*M. rotundifolia* is an ideal partner for breeding PM-resistant grapevines that are durably resistant and require fewer fungicidal applications. This can be facilitated by introgressing functionally diverse PM resistance-associated genes into *V. vinifera* (Michelmore *et al*., 2013). In wild grapes, PM resistance is associated with a programmed cell death-mediated response in infected epidermal cells. This suggests that PM resistance is based on an intracellular recognition of *E. necator*’s effectors by disease resistance (*R*) proteins that activate effector-triggered immunity (Qiu *et al*., 2015; Dry *et al*., 2019).

Most *R* genes encode nucleotide-binding leucine-rich repeat (NLR) proteins (Dubey and Singh, 2018). NLRs are intracellular receptors that recognize and interact directly with pathogen-derived effectors, detect modifications in host cellular targets, or detect molecular decoys triggered by effectors (Dangl *et al*., 2013). NLR activation leads to the induction of immune responses that can restrict pathogen spread (Jones and Dangl, 2006). These include calcium oscillations, a rapid burst of reactive oxygen species, extensive transcriptional reprogramming that leads to cell wall modifications, and the synthesis of pathogenesis-related proteins and antimicrobial compounds (Jones and Dangl, 2006; Dangl *et al*., 2013; Kretschmer *et al*., 2019). Effector-triggered immunity is often associated with a hypersensitive response and programmed cell death of infected plant cells that restrict further pathogen development (Jones and Dangl, 2006). NLR intracellular receptors are typically composed of three domains: a C-terminal LRR domain, a central nucleotide-binding site domain (NBS), and a variable N-terminal domain (Meyers *et al*., 1999; McHale *et al*., 2006). The variable N-terminal domain distinguishes NLR classes. The three main NLR classes are the TIR-NBS-LRRs, CC-NBS-LRRs, and RPW8-NBS-LRRs; these possess N-terminal toll/interleukin-1 receptor-like (TIR), Coiled-coil (CC), and resistance to powdery mildew 8 (RPW8) domains, respectively (Meyers *et al*., 1999; McHale *et al*., 2006; Xiao *et al*., 2001; Michelmore *et al*., 2013). Only two TIR-NBS-LRR genes, *MrRPV1* and *MrRUN1*, have been functionally characterized in grapes (Feechan *et al*., 2013). *MrRPV1* and *MrRUN1* are at the *Run1*/*Rpv1* locus of *M. rotundifolia* G52 and confer resistance to downy mildew and PM, respectively.

The first diploid chromosome-scale genome assembly of a muscadine grape was recently published and represents a valuable resource for identifying candidate PM resistance-associated NLR genes from other genetic loci in *M. rotundifolia* (Cochetel *et al*., 2021). A first analysis of the *Run1*.*2* locus suggested an expansion of TIR-NBS-LRR genes in *M. rotundifolia* Trayshed relative to Cabernet Sauvignon (Cochetel *et al*., 2021). In this study, we describe the structure and gene content of both haplotypes of *Run1*.*2* and *Run2*.*2* loci of *M. rotundifolia* Trayshed. Haplotypes of Trayshed’s *R* loci were differentiated and reconstructed by using deep sequencing data from two backcrossed *V. vinifera* lines, e6-23 (*Run1*.*2b*^+^) and 08391-029 (*Run2*.*2*^+^). Gene models in both loci were manually curated to identify the genes encoding NLRs. The two haplotypes of each *R* locus were compared to determine the effect of heterozygous structural variations on the NLR gene content. To determine NLR genes potentially associated with PM resistance, NLR genes’ expression in *Run1*.*2b* and *Run2*.*2* were profiled with and without PM present using RNA-sequencing (RNA-seq).

## Materials and methods

### Plant material

We used two *V. vinifera* backcross lines in this study: e6-23 carrying the locus *Run1*.*2b* (Feechan *et al*., 2015), and 08391-029 possessing *Run2*.*2* (Riaz *et al*., 2011). Information about the lineage of each genotype is provided in **Supplementary Table S1**. For each grape accession, three plants were inoculated with *E. necator* C-strain and three plants were mock-inoculated as described in Amrine *et al*. (2015). Two leaves from each plant were collected 1 and 5 days post inoculation (dpi) and immediately frozen in liquid nitrogen. Leaves from an individual plant were pooled together and constitute a biological replicate. Three biological replications were obtained for each treatment.

### DNA and RNA extraction, library preparation, and sequencing

Extraction of genomic DNA from mock-inoculated leaves of e6-23 (*Run1*.*2b*^+^) and 08391-029 (*Run2*.*2*^+^) and library preparation were processed as in Massonnet *et al*. (2020). Final libraries were sequenced on the Illumina HiSeqX Ten system in paired-end 150-bp reads (IDseq, Davis, CA, USA) (**Supplementary Table S1**).

RNA extraction and library preparation were performed as in Amrine *et al*. (2015). cDNA libraries were sequenced using Illumina HiSeq2500 and HiSeq4000 sequencers (DNA Technologies Core, University of California, Davis, CA, USA) as 50-bp single-end reads (**Supplementary Table S2**).

### Locus reconstruction

The two haplotypes of *Run1*.*2* and *Run2*.*2* were located by aligning the primers of *Run1*.*2*-associated markers, VMC4f3.1 and VMC8g9, and *Run2*.*2*-associated markers, VMC7f2 and UDV108, onto the diploid chromosome-scale genome of *M. rotundifolia* Trayshed (Riaz *et al*., 2011; Cochetel *et al*., 2021). Whole-genome DNA sequencing reads from e6-23 (*Run1*.*2b*^+^) and 08391-029 (*Run2*.*2*^+^) were then used to identify *Run1*.*2b* and *Run2*.*2* sequences. Low-quality DNA sequencing reads were removed and adapter sequences were trimmed using Trimmomatic v.0.36 (Bolger *et al*., 2014) with the following settings: LEADING:3 TRAILING:3 SLIDINGWINDOW:10:20 MINLEN:36 CROP:150. High-quality paired-end reads were aligned onto the diploid genome of *M. rotundifolia* Trayshed (Cochetel *et al*., 2021) using BWA v.01.17 (Li and Durbin, 2009) and default parameters. Reads aligning onto the reference genome with no edit distance (0 mismatch), were selected using bamtools filter v.2.5.1 (Barnett *et al*., 2011) and the tag “NM:0”. These alignments were used as input for evaluating base coverage with genomecov (BEDTools v2.29.1; Quinlan, 2014). Coverage from bases located in repetitive elements were removed using BEDTools intersect v2.29.1 (Quinlan, 2014). Median coverage per 10 kbp was calculated using an in-house R script and normalized by dividing by the sequencing coverage (**Supplementary Table 1**). Sequences were removed from the locus and labeled “unplaced” if DNA sequencing reads did not cover a primary contig nor its alternative haplotigs. Each haplotype was fragmented into 1-kbp sequences using seqkit sliding v.0.16.1 (Shen *et al*., 2016) and aligned to itself using Minimap2 v.2.12-r847-dirty (Li, 2018). Sequence overlaps between contigs were removed from the locus. DNA sequencing coverage along the four haplotypes was manually inspected by visualizing alignments using Integrative Genomics Viewer (IGV) v.2.4.14 (Robinson *et al*., 2011). Loci were reconstructed using the script HaploMake.py from the tool suite HaploSync v1.0 (https://github.com/andreaminio/HaploSync).

### Haplotype sequence comparison

Pairwise alignments were performed using NUCmer from MUMmer v.4.0.0 (Marçais *et al*., 2018) with the option -- mum. Alignments with at least 90% identity are shown in **Figure 1**. Structural variants (SVs; >50 bp), SNPs and INDELs (<50 bp) were called using show-diff and show-snps, respectively from MUMmer v.4.0.0 (Marçais *et al*., 2018). Potential impact of SNPs on amino acid content was evaluated using SnpEff v.4.3t (Cingolani *et al*., 2012).

**Figure 1:**
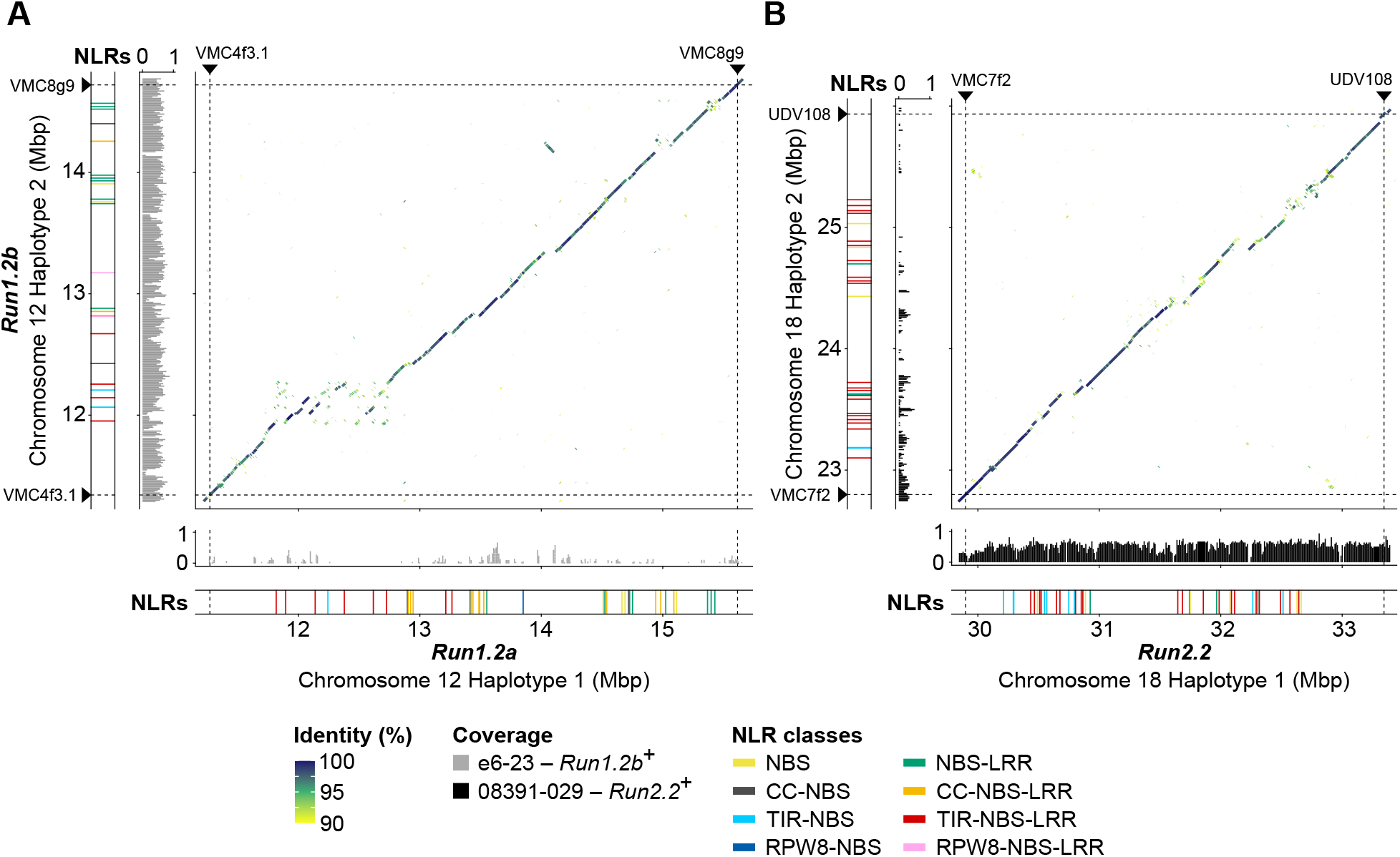
Haplotype comparison and NLR content at *Run1*.*2* and *Run2*.*2* in *M. rotundifolia* Trayshed. Whole-sequence alignments of the reconstructed haplotypes of *Run1*.*2* (**A**) and *Run2*.*2* (**B**) loci. Normalized median DNA-seq coverage per 10 kbp of e6-23 (*Run1*.*2b*^+^) and 08391-029 (*Run2*.*2*^+^) on the diploid genome of *M. rotundifolia* Trayshed was used to identify *Run1*.*2b* and *Run2*.*2* on the Haplotype 2 of chromosome 12 and Haplotype 1 of chromosome 18, respectively. Only DNA-seq reads aligning perfectly on the diploid genome of *M. rotundifolia* Trayshed were used for base coverage analysis. Chromosomal position of the *Run1*.*2*- and *Run2*.*2*-associated genetic markers is indicated by black triangles and dashed lines.

### Annotation of NLR genes

Gene loci potentially associated with NLRs were identified using NLR-annotator with default parameters (Steuernagel *et al*., 2020). Gene models within the *R* loci were manually refined by visualizing alignments of RNA-seq reads from leaves of Trayshed (Cochetel *et al*., 2021), e6-23 (*Run1*.*2b*^+^), and 08391-029 (*Run2*.*2*^+^) using Integrative Genomics Viewer (IGV) v.2.4.14 (Robinson *et al*., 2011). RNA-seq reads were aligned onto the diploid genome of *M. rotundifolia* Trayshed using HISAT2 v.2.1.0 (Kim *et al*., 2015) and the following settings: --end-to-end -- sensitive -k 50.

Predicted proteins were scanned with hmmsearch from HMMER v.3.3.1 (http://hmmer.org/) and the Pfam-A Hidden Markov Models (HMM) database (El-Gebali *et al*., 2019; downloaded on 29 January 2021). Protein domains corresponding to the following Pfam domains: NB-ARC (PF00931.23), LRR (PF00560.34, PF07725.13, PF12799.8, PF13306.7, PF13516.7, PF13855.7), TIR (PF01582.21, PF13676.7), RPW8 (PF05659.12), with an independent E-value less than 1.0, and an alignment covering at least 50% of the HMM were selected (**Supplementary Table S3**). Coiled-coil (CC) domains were identified using COILS (Lupas *et al*., 1991).

### Phylogenetic analysis

Predicted NLR protein sequences from Trayshed’s *Run1*.*2* and *Run2*.*2* and G52’s *Run1/Rpv1* (Feechan *et al*., 2013) were aligned using MUSCLE (Edgar, 2004) in MEGAX (Kumar *et al*., 2018). Resistance gene analogs (RGAs) from *Run1/Rpv1* (Feechan *et al*., 2013) were retrieved on GenBank using the following accession numbers: RGA1, AGC24025; RGA2, AGC24026; RGA4, AGC24027; MrRPV1 (RGA8), AGC24028; RGA9, AGC24029; MrRUN1 (RGA10), AGC24030; RGA11, AGC24031. Phylogenetic analysis of the proteins was performed with MEGAX (Kumar *et al*., 2018) using the Neighbor-Joining method (Saitou and Nei, 1987) and 1,000 replicates.

### Gene expression analysis

Transcript abundance was evaluated with Salmon v.1.5.1 (Patro *et al*., 2017) and the parameters: --gcBias --seqBias --validateMappings. Transcriptome index file was built using a k-mer size of 13 and the combined transcriptomes of *M. rotundifolia* Trayshed, *V. vinifera* cv. Cabernet Sauvignon (Massonnet *et al*., 2020) and *E. necator* C-strain (Jones *et al*., 2014), and their genomes as decoy. Quantification files were imported using the R package tximport v.1.20.0 (Soneson *et al*., 2015). Statistical testing of differential gene expression was performed using DESeq2 v.1.16.1 (Love *et al*., 2014).

## Results

### Structural variants between Trayshed’s *Run1*.*2* haplotypes affect NLR content

The boundaries of *Run1*.*2* were assigned by aligning the primer sequences of *Run1*.*2*-associated SSR markers on the two complete copies (Haplotype 1 & 2) of chromosome 12 of *M. rotundifolia* Trayshed (Cochetel *et al*., 2021). To distinguish *Run1*.*2a* and *Run1*.*2b*, we sequenced the genome of the *V. vinifera* backcross e6-23 (*Run1*.*2b*^+^), into which *Run1*.*2b* was introgressed by crossing with *M. rotundifolia* Trayshed and backcrossing with *V. vinifera* (**Supplementary Tables S1 & S4**). Short sequencing reads from the *Run1*.*2b*^+^ accession covered and aligned perfectly (*i*.*e*., with no mismatches) to most of *Run1*.*2* on chromosome 12 Haplotype 2 (**Supplementary Figure S1**), and coverage gaps in *Run1*.*2* on Haplotype 2 were complemented by coverage at *Run1*.*2* on Haplotype 1. This indicates that haplotype switching occurred during the assembly and phasing of Trayshed’s genome. To correct this, *Run1*.*2b* was reconstructed using only sequences supported with DNA sequencing reads from the *Run1*.*2b*^+^ accession and *Run1*.*2a* was reconstructed using alternative sequences (**Figure 1A**). The two reconstructed *Run1*.*2* haplotypes, *Run1*.*2a* and *Run1*.*2b*, were 4.34 Mbp-and 3.38 Mbp-long, respectively. Differences in length between the two haplotypes were associated with several large structural variants (SVs; > 50 bp). For instance, the region of *Run1*.*2b* from ∼12 Mbp to 12.3 Mbp corresponds to a ∼800 kbp region in the *Run1*.*2a* haplotype (**Figure 1A**). In this case, length difference was due to several inserted sequences and duplication events in *Run1*.*2a* compared to *Run1*.*2b*. Furthermore, we detected 32,704 SNPs and 7,150 INDELs between *Run1*.*2a* and *Run1*.*2b*.

To determine the effect of the heterozygous SVs and short polymorphisms on the gene content, we refined the gene models for both *Run1*.*2* haplotypes. A total of 78 protein-coding genes, including 22 NLR genes, were manually annotated (**Table 1**). *Run1*.*2a* contained 253 genes and *Run1*.*2b* contained 189 genes, indicating that SVs affect the gene content. There were 37 and 24 NLR genes in *Run1*.*2a* and *Run1*.*2b*, respectively, with both composed primarily of CC-NBS-LRR, TIR-NBS-LRR and NBS-LRR genes (**Figure 1A**; **Supplementary Table S4**). SVs between haplotypes affect the protein-coding sequences of 22 NLR genes in *Run1*.*2a* and nine NLR genes in *Run1*.*2b*. These SVs resulted in the whole duplication of four and two NLR genes in *Run1*.*2a* and *Run1*.*2b*, respectively, and the partial duplication of three NLR genes in *Run1*.*2a* (**Supplementary Table S4**). In addition, SVs were found to cause the loss of functionality of four NLR-coding genes and the hemizygosity of a CC-NBS gene in *Run1*.*2a* relative to *Run1*.*2b*, as well as the loss of the LRR domain of two NLR genes in *Run1*.*2b* compared to *Run1*.*2a*. In addition, we detected 32,704 SNPs and 7,150 INDELs between the two *Run1*.*2* haplotypes (*Run1*.*2*a *vs. Run1*.*2b*). Non-synonymous SNPs were identified in eight NLR genes in each haplotype.

**Table 1:** Sequence length, protein-coding gene content, and NLR gene content of *Run1*.*2* and *Run2*.*2* reconstructed haplotypes. Numbers in parentheses correspond to the genes with a structure manually refined.

| **Loci** | ***Run1.2a*** | ***Run1.2b*** | ***Run2.2*** | **Chr18 Hap2** |
| --- | --- | --- | --- | --- |
| Sequence length (bp) | 4,340,059 | 3,379,591 | 3,445,914 | 3,137,389 |
| Protein-coding gene loci | 253 (44) | 189 (34) | 207 (59) | 179 (43) |
| Total NLR genes | 37 (16) | 24 (6) | 39 (33) | 29 (22) |
| NBS genes | 3 (1) | 2 | 6 (6) | 3 (3) |
| CC-NBS genes | 2 (1) | 2 | 0 | 0 |
| RPW8-NBS genes | 2 (2) | 0 | 0 | 0 |
| TIR-NBS genes | 1 (1) | 2 (1) | 9 (7) | 3 (1) |
| NBS-LRR genes | 8 (2) | 9 | 4 (4) | 2 (2) |
| CC-NBS-LRR genes | 13 (6) | 3 (1) | 0 | 0 |
| RPW8-NBS-LRR genes | 0 | 2 | 0 | 0 |
| TIR-NBS-LRR genes | 8 (3) | 4 (4) | 20 (16) | 21 (16) |

### *Run2*.*2* is mainly composed of TIR-NBS-LRRs

A similar approach was applied to identify and reconstruct *Run2*.*2* in the Haplotype 1 of chromosome 18 of *M. rotundifolia* Trayshed using short sequencing reads from the genotype 08391-029 (*Run2*.*2*^+^) (**Supplementary Tables S1 & S4**; **Supplementary Figure S2**). The reconstructed *Run2*.*2* was 3.45 Mbp long, slightly longer than its alternative on Haplotype 2 (3.14 Mbp). We manually refined the models of 102 protein-coding genes in the two haplotypes, including 55 NLR genes (**Table 1**). More genes were annotated at *Run2*.*2* (207) than at its alternative (179). There were 39 NLR-coding genes at *Run2*.*2* and 29 NLR genes in its alternative. The two haplotypes were mainly composed of TIR-NBS-LRR genes, with 20 genes in *Run2*.*2* and 21 genes in its alternative (**Supplementary Table S4**). Unlike *Run1*.*2*, no NLR genes with a CC or RPW8 N-terminal domain were found at *Run2*.*2*. Interestingly, the NLR genes occurred in two clusters in each haplotype (**Figure 1B**; **Table 1**).

*Run2*.*2* and its alternative contained 456 SVs between them, with an average length of 2.1 ± 3.6 kbp. These SVs affected 21 and 16 NLR genes in *Run2*.*2* and its alternative, respectively. SVs were found responsible for the partial duplication of four and two NLR genes in *Run2*.*2* and its alternative haplotype, respectively (**Supplementary Table S4**). Furthermore, large deletions encompassed the complete coding sequence of three and four NLR-coding genes of *Run2*.*2* and its alternative haplotype, respectively. We also identified 24,128 SNPs and 5,773 INDELs between *Run2*.*2* and its alternative, and non-synonymous SNPs were detected in 16 and 18 NLR genes, respectively.

### *Run1*.*2* and *Run2*.*2* loci contain distinct sets of NLRs

Predicted protein sequences of the NLRs identified in the four reconstructed haplotypes: *Run1*.*2a, Run1*.*2b, Run2*.*2* and its alternative on chromosome 18 Haplotype 2, were compared by constructing a phylogenic tree (**Figure 2A**). In addition, we compared Trayshed’s NLRs with the TIR-NBS-LRRs at *Run1/Rpv1* in *M. rotundifolia* G52 (Barker *et al*., 2005; Feechan *et al*., 2013). *Run1/Rpv1* is the only *R* locus characterized in grapes and is an alternative version of *Run1*.*2*. Two distinct groups of NLRs were discovered, distinguished by the presence or absence of a TIR domain. A similar clustering pattern was observed when a phylogeny was built using NBS domain sequences only (**Supplementary Figure S3**), as previously observed in other plants (Prigozhin and Krasileva, 2021; Seo *et al*., 2016). NLRs also tended to cluster by *R* locus, indicating allele relationship between haplotypes for 74.4% of the NLRs (**Supplementary Table S4**). Regarding the TIR-containing proteins, we found the TIR-NBS-LRRs from *Run1/Rpv1* clustering with the TIR-NBS-LRR proteins from *Run1*.*2*. MrRPV1 from *M. rotundifolia* G52 clustered with two TIR-NBS-LRRs, one from each *Run1*.*2* haplotype, and MrRUN1 clustered with a TIR-NBS-LRR from *Run1*.*2a* (**Figure 2A**; **Supplementary Table S4**). Clustering of TIR-NBS-LRRs of *Run1*.*2* and *Run1/Rpv1* support an allelic relationship between them. However, number of LRR motifs in their LRR domain was different (**Figure 2B**), suggesting some allelic diversity. In addition, differences in LRR domains suggest that these TIR-NBS-LRRs might be specific to different effectors and/or pathogens (McHale *et al*., 2006).

**Figure 2:**
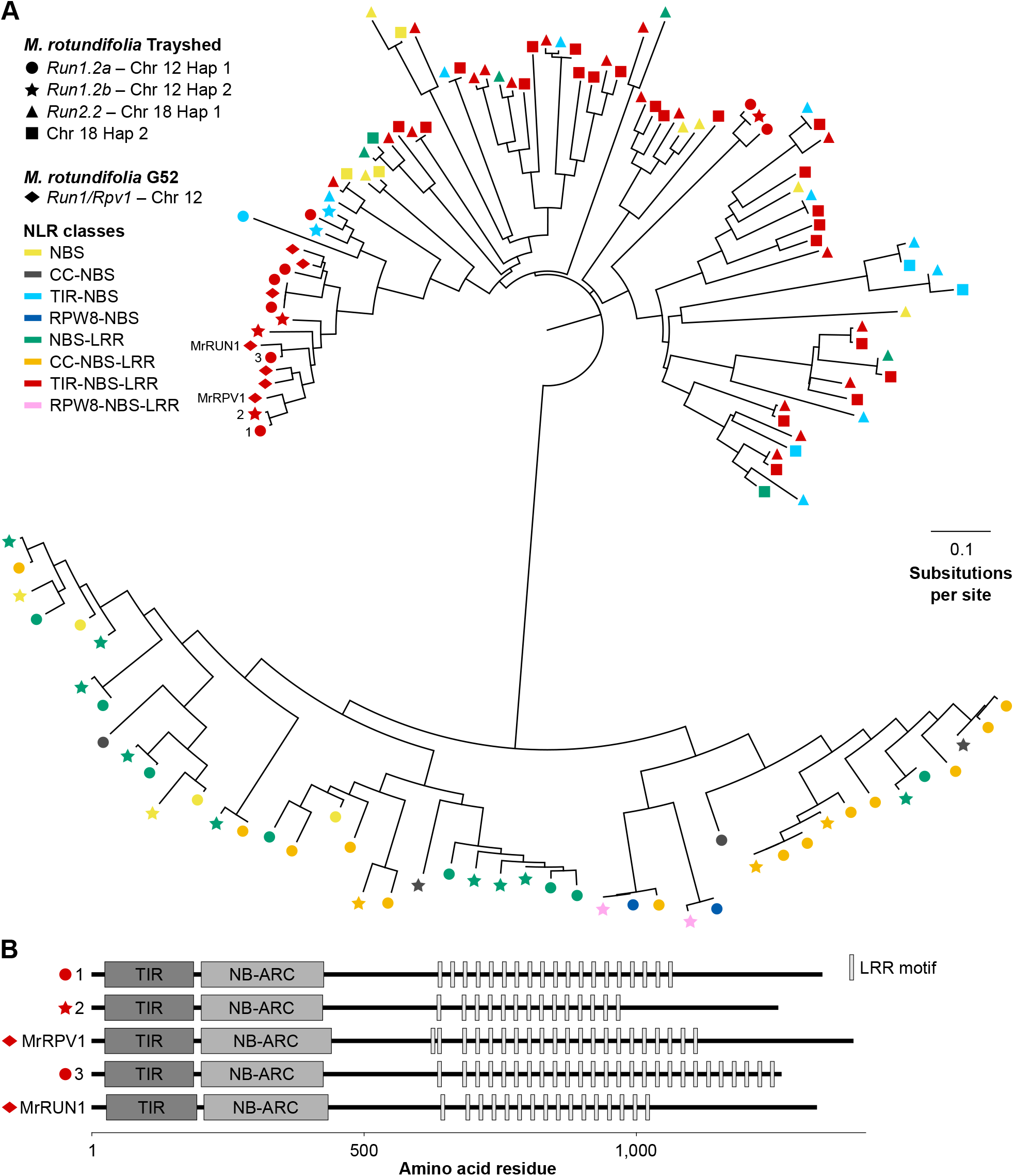
Comparison of the NLRs composing *Run1*.*2* and *Run2*.*2* in *M. rotundifolia* Trayshed. (**A**) Neighbor-joining clustering of the predicted protein sequences of the NLRs composing *Run1*.*2* and *Run2*.*2* haplotypes, and *Run1/Rpv1* (Feechan *et al*., 2013). (**B**) Domain diagram of Trayshed’s TIR-NBS-LRRs clustering with G52’s MrRUN1 and MrRPV1. Proteins are reported using same number assigned in panel **A**. LRR motifs were identified using the consensus sequence LxxLxLxx, with L indicating a leucine residue and x indicating any amino acid (Kajava and Kobe, 2002).

### Most of NLR genes at *Run1*.*2b* and *Run2*.*2* are constitutively expressed

Constitutive NLR gene expression is essential for disease resistance (Michelmore *et al*., 2013). To identify expressed NLR genes that are potentially responsible for PM resistance, we measured gene expression in *Run1*.*2b*^+^ and *Run2*.*2*^+^ leaves 1 and 5 days after inoculation (dpi) with either *E. necator* C-strain or a mock solution using RNA-seq (**Figure 3**).

**Figure 3:**
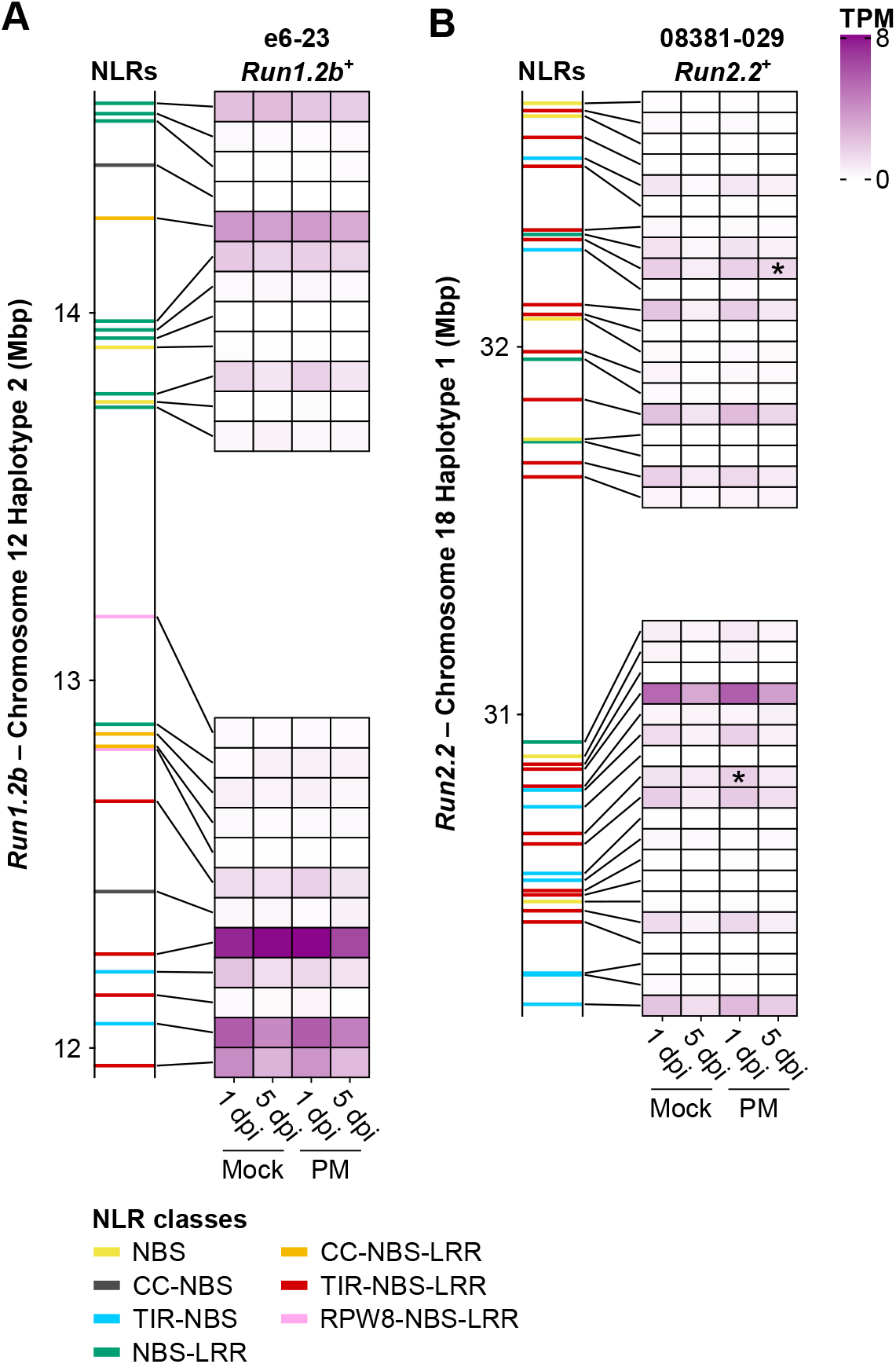
Transcript abundance of NLR genes in *Run1*.*2b* (**A**) and *Run2*.*2* (**B**). Gene expression was monitored in *Run1*.*2b*^+^ and *Run2*.*2*^+^ at 1 and 5 days post inoculation (dpi) with *E. necator* conidia (PM) or a mock solution (Mock). Transcript abundance is shown as the mean of Transcripts per Million (TPM). n = 3. NLR genes differentially expressed in response to PM are indicated by an asterisk (*P* ≤ 0.05).

In *Run1*.*2b*^+^ leaves, expression of almost all NLR genes from the *Run1*.*2b* locus (23/24) was detected (**Figure 3A**). Nine of them were found with an expression level higher than 1 transcript per million (TPM) in at least one condition (TPM > 1), while 15 NLR genes exhibit a lower expression level (TPM ≤ 1). Three genes at the 5’-end of *Run1*.*2b* encoding two TIR-NBS-LRRs and a TIR-NBS, and a CC-NBS-LRR gene towards the 3’-end were the most highly expressed in all conditions (mean TPM > 4 TPM). In addition, the gene with the most elevated expression was the TIR-NBS-LRR gene which predicted protein clustered with MrRPV1 in the phylogenic tree (**Figure 2**; **Supplementary Table S4**). PM inoculation was not found to significantly impact on the gene expression of any NLR gene composing *Run1*.*2b*.

In *Run2*.*2*^+^ genotype, we identified 11 NLR genes with a transcript abundance superior to 1 TPM, 24 lowly expressed (TPM ≤ 1) and four with no expression (**Figure 3B**). A TIR-NBS-LRR gene at the 5’-end of *Run2*.*2* was the most highly expressed across conditions. Seven other TIR-NBS-LRR genes in the locus had moderate expression levels. Only two NLR genes of the *Run2*.*2* locus were modulated in response to PM, one at 1 dpi and another one at 5 dpi.

These RNA-seq data show that most of the NLR genes composing Trayshed’s loci are expressed independently of the condition, although they exhibit different levels of gene expression in *Run1*.*2b*^+^ and *Run2*.*2*^+^ genotypes. Expressed NLR genes are candidate genes involved in PM resistance associated with *Run1*.*2b* and *Run2*.*2*. All candidate genes, their coordinates, expression, and relation to known *Run1/Rpv1*-associated NLRs are reported in **Supplementary Table S4**.

## Discussion

By combining a diploid assembly of Trayshed’s genome and DNA sequencing data generated from two backcrossed *V. vinifera* genotypes, we distinguished, phased, and reconstructed the four complete haplotypes of *Run1*.*2* and *Run2*.*2* loci. To our knowledge, this is the first report of complete, haplotype-resolved *R* loci for grape. The *Run1*/*Rpv1* locus was sequenced prior, but its assembly was fragmented and haploid (Feechan *et al*., 2013). Trayshed was previously defined as homozygous at *Run2*.*2 locus* based on amplicon size (Riaz *et al*., 2011). However, *in silico* PCR on Trayshed’s genome showed two amplicon sizes for the UDV108 marker: 225 bp on chromosome 18 Haplotype 1 and 323 bp on chromosome 18 Haplotype 2 (**Supplementary Table S4**). Sequencing additional *Run2*.*2*^+^ backcrossed individuals would help determine whether the region on chromosome 18 Haplotype 2 is associated with PM resistance.

Manual annotation was performed for 77 NLR genes, representing ∼60% of the NLR genes identified among the four haplotypes (**Table 1**). Combined with haplotype resolution, it allowed the characterization of NLR gene variation between haplotypes of each locus. This highlights the necessity of generating phased haplotypes and meticulously dissecting complex genomic regions if candidate trait-associated genes are sought (Massonnet *et al*., 2020).

The NLRs in Trayshed’s *Run1*.*2* and *Run2*.*2* loci differ. All three classes of NLRs, CC-NBS-LRRs, RPW8-NBS-LRRs and TIR-NBS-LRRs were found in *Run1*.*2* haplotypes, but no CC or RPW8 domains were identified among NLRs at *Run2*.*2*. The only characterized NLR gene associated with grape PM resistance, MrRUN1, is a TIR-NBS-LRR (Feechan *et al*., 2013). Functional characterization of the NLR genes composing Trayshed’s *R* loci would help identify the NLR(s) responsible for PM resistance and to determine whether the NLR class is a decisive factor for grape PM resistance.

Most of the NLR genes composing *Run1*.*2b* and *Run2*.*2* were found expressed at low level independently of the presence of the pathogen. Constitutive low expression of NLR genes is commonly found in plants (Michelmore *et al*., 2013; Zhang *et al*., 2020), supporting a constitutive ability to sense pathogens. However, some plant NLR genes were found to be more highly expressed during pathogen infection, suggesting an induction of the defense-related surveillance in response to biotic stress (Mohr *et al*., 2010; Sagi *et al*., 2017; Zhang *et al*., 2020). Only three NLR genes were found differentially expressed in response to *E. necator* C-strain in the two backcrossed *V. vinifera* genotypes possessing *Run1*.*2b* and *Run2*.*2*. It would be interesting to assess gene expression level and transcriptional modulation of the NLR genes composing *Run1*.*2a* with the same protocol, as well as using different individuals carrying Trayshed’s PM-associated loci to evaluate the effect of the genetic background on NLR gene expression. In addition, monitoring NLR gene expression in response to additional *E. necator* strains would help determining whether the NLR genes composing the two PM-associated loci exhibit a strain-specific gene expression and/or transcriptional modulation.

Finally, the approach used in this study could be applied to other *R* loci, giving new opportunities for functional genomics. However, fine mapping and generation of sequencing data (DNA-seq and RNA-seq) from recombinants would be necessary to narrow down candidate NLR genes associated with PM resistance.

## Data availability

Sequencing data are accessible through NCBI under the BioProject ID PRJNA780568. New genome assembly of *M. rotundifolia* Trayshed and annotation files are available at Zenodo under the DOI 10.5281/zenodo.5703495 and at www.grapegenomics.com.

## Supplementary information

**Supplementary Figure S1**: Reconstruction of *Run1*.*2a* and *Run1*.*2b* loci in *M. rotundifolia* Trayshed. Self-comparison of *Run1*.*2a* (chromosome 12 Haplotype 1) and *Run1*.*2b* (chromosome 12 Haplotype 2) loci, normalized median coverage of DNA-seq reads from e6-23 (*Run1*.*2b*^+^) and Trayshed (*Run1*.*2a/b*^+^) before (**A**) and after (**B**) reconstruction. Only DNA-seq reads aligning perfectly on the diploid genome of *M. rotundifolia* Trayshed were used for the base coverage analysis. Chromosomal position of the genetic markers VMC4f3.1 and WMC8g9 is indicated by black triangles and dashed lines.

**Supplementary Figure S2**: Reconstruction of *Run2*.*2* locus and its alternative haplotype in *M. rotundifolia* Trayshed. Self-comparison of *Run2*.*2* (chromosome 18 Haplotype 1) and its alternative haplotype (chromosome 18 Haplotype 2) loci, normalized median coverage of DNA-seq reads from 08391-029 (*Run2*.*2*^+^) and Trayshed (*Run2*.*2*^+^) before (**A**) and after (**B**) reconstruction. Only DNA-seq reads aligning perfectly on the diploid genome of *M. rotundifolia* Trayshed were used for the base coverage analysis. Chromosomal position of the genetic markers VMC7f2 and UDV108 is indicated by black triangles and dashed lines.

**Supplementary Figure S3**: Neighbor-joining clustering of the nucleotide-binding domain of the predicted NLR protein sequences of the loci *Run1*.*2, Run2*.*2* and *Run1*/*Rpv1* (Feechan *et al*., 2013). **Supplementary Table S1**: Pedigree information of e6-23 (*Run1*.*2b*^+^) and 08391-029 (*Run2*.*2*^+^) and summary statistics of DNA sequencing.

**Supplementary Table S2**: Summary statistics of RNA sequencing.

**Supplementary Table S3**: Domain composition of the NLRs identified at *Run1*.*2* and *Run2*.*2*. **Supplementary Table S4**: Summary of the NLRs composing *Run1*.*2* and *Run2*.*2* loci. This table includes the chromosomal position of *Run1*.*2*- and *Run2*.*2*-associated SSR markers, as well as the chromosomal position of the NLR genes, the NLR class of the predicted proteins, the effect of the SVs detected between haplotypes, the NLR protein clustering in the phylogenetic tree represented in Figure 2A, and their average gene expression (TPM) in leaves of e6-23 (*Run1*.*2b*^+^) and 08391- 029 (*Run2*.*2*^+^).

## Literature cited

Amrine, K. et al. Comparative transcriptomics of Central Asian Vitis vinifera accessions reveals distinct defense strategies against powdery mildew. Hortic Res 2, 15037 (2015).

Barker, C. L. et al. Genetic and physical mapping of the grapevine powdery mildew resistance gene, Run1, using a bacterial artificial chromosome library. Theor. Appl. Genet. 111, 370–377 (2005).

Barnett, D. W., Garrison, E. K., Quinlan, A. R., Strömberg, M. P. & Marth, G. T. BamTools: a C++ API and toolkit for analyzing and managing BAM files. Bioinformatics 27(12), 1691–1692 (2011).

Bolger, A. M., Lohse, M. & Usadel, B. Trimmomatic: a flexible trimmer for Illumina sequence data. Bioinformatics 30, 2114–2120 (2014).

Cingolani, P. et al. A program for annotating and predicting the effects of single nucleotide polymorphisms, SnpEff: SNPs in the genome of Drosophila melanogaster strain w1118; iso-2; iso-3. Fly 6(2), 80–92 (2012).

Cochetel, N. et al. Diploid chromosome-scale assembly of the Muscadinia rotundifolia genome supports chromosome fusion and disease resistance gene expansion during Vitis and Muscadinia divergence. G3 (Bethesda, Md.) 11(4), jkab033 (2021).

Dalbó, M. A., Ye, G. N., Weeden, N. F., Wilcox, W. F. & Reisch, B. I. Marker-assisted selection for powdery mildew resistance in grapes. J Amer Soc Hort Sci 126(1), 83–89 (2001).

Dangl, J. L., Horvath, D. M. & Staskawicz, B. J. Pivoting the plant immune system from dissection to deployment. Science 341(6147), 746–751 (2013).

Dry, I., Riaz, S., Fuchs, M., Sosnowski M. & Thomas, M. Scion breeding for resistance to biotic stresses. In The Grape Genome (eds Cantu, D. & Walker, M. A) Ch. 15 (Spinger, 2019).

Dubey, N. & Singh, K. Role of NBS-LRR proteins in plant defense. In Molecular Aspects of Plant-Pathogen Interaction (eds Singh A. & Singh I.) (Springer, 2018).

Edgar, R. C. MUSCLE: multiple sequence alignment with high accuracy and high throughput. Nucleic Acids Res. 32, 1792–1797 (2004).

El-Gebali, S. J. et al. The Pfam protein families database in 2019. Nucleic Acids Res. 47, D427–D432 (2019).

Feechan, A. et al. Genetic dissection of a TIR-NB-LRR locus from the wild North American grapevine species Muscadinia rotundifolia identifies paralogous genes conferring resistance to major fungal and oomycete pathogens in cultivated grapevine. Plant J. 76, 661–674 (2013).

Feechan, A. et al. Strategies for RUN1 deployment using RUN2 and REN2 to manage grapevine powdery mildew informed by studies of race specificity. Phytopathology 105(8), 1104–1113 (2015).

Gadoury, D. M. et al. Grapevine powdery mildew (Erysiphe necator): a fascinating system for the study of the biology, ecology and epidemiology of an obligate biotroph. Mol Plant Pathol. 13, 1–16 (2012).

Jones, J. & Dangl, J. The plant immune system. Nature 444, 323–329 (2006).

Jones, L. et al. Adaptive genomic structural variation in the grape powdery mildew pathogen, Erysiphe necator. BMC Genomics 15(1), 1081 (2014).

Kajava, A. V. & Kobe, B. Assessment of the ability to model proteins with leucine-rich repeats in light of the latest structural information. Protein Sci 11, 1082–1090 (2002).

Karn, A. et al. Discovery of the REN11 locus from Vitis aestivalis for stable resistance to grapevine powdery mildew in a family segregating for several unstable and tissue-specific quantitative resistance loci. Front Plant Sci. 12, 733899 (2021).

Kim, D., Langmead, B. & Salzberg, S. L. HISAT: a fast spliced aligner with low memory requirements. Nat. Methods 12, 357–360 (2015).

Kretschmer, M., Damoo, D., Djamei, A. & Kronstad, J. Chloroplasts and plant immunity: Where are the fungal effectors? Pathogens 9(1), 19 (2019).

Kumar, S., Stecher, G., Li, M., Knyaz, C. & Tamura, K. MEGA X: Molecular Evolutionary Genetics Analysis across computing platforms. Mol Biol Evol. 35, 1547–1549 (2018).

Li, H. & Durbin, R. Fast and accurate short read alignment with Burrows−Wheeler transform. Bioinformatics 25(14), 1754–1760 (2009).

Li, H. Minimap2: pairwise alignment for nucleotide sequences. Bioinformatics 34(18), 3094–3100 (2018).

Love, M. I., Huber, W. & Anders, S. Moderated estimation of fold change and dispersion for RNA-seq data with DESeq2. Genome Biol 15, 550 (2014).

Lupas, A., Van Dyke, M. & Stock, J. Predicting coiled coils from protein sequences. Science 252, 1162–1164 (1991).

Marçais, G. et al. MUMmer4: a fast and versatile genome alignment system. PLoS Comput. Biol. 14, e1005944 (2018).

Massonnet, M. et al. The genetic basis of sex determination in grapes. Nat Commun 11, 2902 (2020).

McHale, L., Tan, X., Koehl, P. & Michelmore, R. W. Plant NBS-LRR proteins: adaptable guards. Genome Biol 7, 212 (2006).

Meyers, B. C. et al. Plant disease resistance genes encode members of an ancient and diverse protein family within the nucleotide binding superfamily. Plant J 20, 317–332 (1999).

Michelmore, R. W., Christopoulou, M. & Caldwell, K. S. Impacts of resistance gene genetics, function, and evolution on a durable future. Annu Rev Phytopathol. 51, 291–319 (2013).

Mohr, T. J. et al. The Arabidopsis downy mildew resistance gene RPP8 is induced by pathogens and salicylic acid and is regulated by Wbox cis elements. Mol. Plant-Microbe Interact. 23, 1303–1315 (2010).

Olmo, H. P. Vinifera rotundifolia hybrids as wine grapes. Am. J. Enol. Vitic. 22, 87–91 (1971).

Olmo, H. P. The potential role of (vinifera x rotundifolia) hybrids in grape variety improvement. Experientia 42, 921–926 (1986).

Pap, D. et al. Identification of two novel powdery mildew resistance loci, Ren6 and Ren7, from the wild Chinese grape species Vitis piasezkii. BMC Plant Biol. 16, 170 (2016).

Patro, R., Duggal, G., Love, M. I., Irizarry, R. A. & Kingsford, C. Salmon provides fast and bias-aware quantification of transcript expression. Nat Methods 14(4), 417–419 (2017).

Pauquet, J. et al. Establishment of a local map of AFLP markers around the powdery mildew resistance gene Run1 in grapevine and assessment of their usefulness for marker assisted selection. Theor Appl Genet 103, 1201–1210 (2001).

Prigozhin, D. M. & Krasileva, K. V. Analysis of intraspecies diversity reveals a subset of highly variable plant immune receptors and predicts their binding sites. Plant Cell 33(4), 998–1015 (2021).

Qiu, W., Feechan, A. & Dry, I. Current understanding of grapevine defense mechanisms against the biotrophic fungus (Erysiphe necator), the causal agent of powdery mildew disease. Hortic Res 2, 15020 (2015).

Quinlan, A. R. BEDTools: the Swiss-army tool for genome feature analysis. Curr Protoc Bioinformatics 47, 11.12.1-34 (2014).

Ramming, D. W. et al. A single dominant locus, Ren4, confers rapid non-race-specific resistance to grapevine powdery mildew. Phytopathology 101, 502–508 (2011).

Riaz, S., Tenscher, A. C., Ramming, D. W. & Walker, M. A. Using a limited mapping strategy to identify major QTLs for resistance to grapevine powdery mildew (Erysiphe necator) and their use in marker-assisted breeding. Theor Appl Genet 122, 1059–1073 (2011).

Riaz, S., Menéndez, C. M., Tenscher, A., Pap, D. & Walker, M. A. Genetic mapping and survey of powdery mildew resistance in the wild Central Asian ancestor of cultivated grapevines in Central Asia. Hort Res 7, 104 (2020).

Robinson, J. T. et al. Integrative genomics viewer. Nat. Biotechnol. 29, 24–26 (2011).

Sagi, M. S., Deokar, A. A. & Tar’an, B. Genetic analysis of NBS-LRR gene family in chickpea and their expression profiles in response to Ascochyta blight infection. Front Plant Sci., 8 838 (2017).

Saitou, N. & Nei, M. The neighbor-joining method: A new method for reconstructing phylogenetic trees. Mol. Biol. Evol. 4, 406–425 (1987).

Sambucci, O., Alston, J. M., Fuller, K. B. & Lusk, J. The pecuniary and non-pecuniary costs of powdery mildew, and the potential value of resistant varieties in California grapes. Am J Enol Vitic 70(2), 177–187 (2019).

Seo, E., Kim, S., Yeom, S. I. & Choi, D. Genome-wide comparative analyses reveal the dynamic evolution of nucleotide-binding leucine-rich repeat gene family among Solanaceae plants. Front Plant Sci. 7, 1205 (2016).

Shen, W., Le, S., Li, Y. & Hu, F. SeqKit: A cross-platform and ultrafast toolkit for FASTA/Q file manipulation. PloS One 11(10), e0163962 (2016).

Small, J. K. (eds) Flora of the southeastern United States: being descriptions of the seed-plants, ferns and fern-allies growing naturally in North Carolina, South Carolina, Georgia, Florida, Tennessee, Alabama, Mississippi, Arkansas, Louisiana and in Oklahoma and Texas east of the one hundredth meridian. (J.K. Small, 1913).

Soneson, C., Love, M. I. & Robinson, M. D. Differential analyses for RNA-seq: transcript-level estimates improve gene-level inferences. F1000Res. 4, 1521 (2015).

Steuernagel, B. K. et al. The NLR-Annotator tool enables annotation of the intracellular immune receptor repertoire. Plant Physiol. 183, 468–482 (2020).

Xiao, S. et al. Broad spectrum mildew resistance in Arabidopsis thaliana mediated by RPW8. Science 291, 118–120 (2001).

Zhang, Y. M. et al. Genome-wide identification and evolutionary analysis of NBS-LRR genes from Dioscorea rotundata. Frontiers in genetics 11, 484 (2020).

